# Xylem sap residue in cut-open conduits can affect gas discharge in pneumatic experiments

**DOI:** 10.1101/2023.08.08.552466

**Authors:** Marcela T. Miranda, Luciano Pereira, Gabriel S. Pires, Xinyi Guan, Luciano M. Silva, Swetlana Kreinert, Eduardo C. Machado, Steven Jansen, Rafael V. Ribeiro

**Affiliations:** Laboratory of Plant Physiology ‘Coaracy M. Franco’, Center for Agricultural and Post-Harvest Biosystems, Agronomic Institute (IAC), Campinas, SP, Brazil; Institute of Botany, Ulm University, Albert-Einstein-Allee 11, 89081 Ulm, Germany; Laboratory of Crop Physiology (LCroP), Department of Plant Biology, Institute of Biology, University of Campinas (UNICAMP), Campinas, SP, Brazil

**Keywords:** Embolism, Vulnerability curves, Pneumatron, artefact, *Citrus*

## Abstract

- Considerable progress has been made in understanding the mechanisms of embolism formation based on the pneumatic method, which relies on gas discharge measurements. Here, we test the assumption that cut-open conduits are gas-filled when samples are cut at high water potentials.
- We performed vulnerability curves (VC) with the Pneumatron and analysed sap extraction from cut-open vessels in *Citrus* branches, while the optical method was applied as a reference method. VCs of 11 additional angiosperms were analysed to generalise our findings.
- We found an increase in gas discharge during early stages of dehydration, which affected the VC of *Citrus*. Xylem sap was not absorbed immediately by surrounding tissue in cut *Citrus* branches. The gas amount discharged increased until all sap residue was absorbed, which was near the turgor loss point. By analysing the slope of VCs, we could correct pneumatic VC, as evidenced by the strong agreement in embolism resistance between the pneumatic and the optical method.
- Since residual sap in cut-open conduits of some species could slightly reduce embolism resistance in some species, we propose to apply an easy correction for this novel artefact. Automated measurements with a Pneumatron are also required because of its high time resolution.

## Introduction

Xylem sap in water conducting cells of plants is frequently under negative pressure, and therefore prone to changes in liquid-gas interfaces, which may lead to transport failure by embolism (Dixon & Joly, 1895). Studies on the mechanisms behind sap transport, as well as embolism formation, are hampered by size dimensions of the xylem pathway, varying from micrometre wide conduits to nanometre sized pore constrictions in pit membranes, and the dynamic complexity of multiphase interactions between liquids, gasses, and cell walls (Kaack *et al*., 2021). Moreover, manipulation of xylem tissue, especially when sap is under negative pressure, can result in artefacts when evaluating plant embolism (Wheeler *et al*., 2013; Jansen & Schenk, 2015; Lamarque *et al*., 2018).

Embolism resistance varies among species and is typically evaluated by a vulnerability curve (VC), which gives the relationship between xylem water potential and the corresponding degree of xylem embolism (Pereira *et al*., 2016). When a plant dehydrates, its xylem water potential decreases, and the likelihood of embolism formation may increase. Based on VCs, it is possible to estimate the water potentials inducing 12%, 50% and 88% of embolism (Ψ_12_, Ψ_50_ and Ψ_88_ respectively). While Ψ_12_ is considered to be the threshold for air entry in conduits causing embolism, Ψ_50_ is commonly used to compare the characteristic embolism resistance of a particular species to embolism, and Ψ_88_ represents the water potential associated with the threshold leading to irreversible drought damage and mortality risk (Urli *et al*., 2013).

Several methods have been developed to study xylem vulnerability to embolism, which may be estimated either indirectly by measuring the loss of conductivity due to embolism formation, or directly by quantifying the gas volume of embolised vessels (Venturas *et al*., 2019). The loss of conductivity can be measured with a hydraulic apparatus (Sperry *et al*., 1988) or by the centrifuge method (Cochard, 2002; Peng *et al*., 2019), whereas embolism can be quantified through imaging methods (Brodersen *et al*., 2010; Brodribb *et al*., 2016; Meixner *et al*., 2020), or gas extraction using the Pneumatic method (Pereira *et al*., 2016, 2020).

The pneumatic method quantifies the amount of gas that can be extracted from intact (i.e. non-cut-open) and embolised conduits that are connected via interconduit pit membranes to cut-open conduits. Increases in the discharged air volume during progressive dehydration are then related to the progressive spread of embolism in several series of intact conduits (Pereira *et al*., 2016). When embolism has been induced in intact conduits, a larger gas volume is extracted than prior to embolism propagation (Guan *et al*., 2021; Yang *et al*., 2023). Recently, Pneumatron devices have been used to obtain pneumatic vulnerability curves (Pereira *et al*., 2020). These devices are automated instruments that capture at a high temporal resolution the amount of gas extracted from xylem conduits and therefore evaluate the gas kinetics during xylem dehydration over several hours to days. Detailed modelling of gas diffusion kinetics that underlies pneumatic vulnerability curves, combined with a comparison of various methods, provide strong evidence to consider the pneumatic method as a highly accurate, easy, and low-budget method to estimate embolism resistance of xylem (Guan *et al*., 2021; Brum *et al*., 2023; Yang *et al*., 2023; Paligi *et al.,* In Press).

Nevertheless, various users have already shown that pneumatic measurements should be interpreted carefully to avoid misinterpretation of data, considering also basic assumptions (e.g., Chen *et al*., 2021; Brum *et al*., 2023). A central assumption, for instance, is that conduits cut open in air under atmospheric conditions become fully gas-filled because sap in cut-open conduits would be quickly withdrawn by sap under negative pressure in neighbouring, intact conduits (Van Ieperen *et al*., 2001; Tyree & Zimmermann, 2002). During preliminary work on pneumatic VCs of *Citrus*, however, we suspected that this assumption may not be correct, and that sap in cut-open conduits could have consequences for the amount of gas discharged in pneumatic experiments. If sap would remain in cut-open conduits, this may especially affect gas discharge measurements during initial dehydration stages, which could undermine the stability of the minimum amount of gas discharged prior to embolism propagation. It is important that cut-open conduits in pneumatic experiments are filled with gas because these conduits function as an extension of the discharge tube of the Pneumatron.

Here, we tested if cut-open conduits are fully filled with gas, and whether partial or complete sap removal affected estimations of embolism resistance in pneumatic experiments. Besides *Citrus*, we aimed to examine pneumatic vulnerability curves of 11 additional angiosperm species to evaluate potential problems during high xylem water potentials, and how pneumatic measurements should be analysed and interpreted. We also examined if sap removal from the cut-open vessels is related to the turgor loss point (Ψ_TLP_), as xylem water potentials below the turgor loss point should cause complete absorption of xylem sap from the cut-open vessels by its surrounding tissue.

## Material and Methods

### Plant material

We constructed vulnerability curves (VCs) using the Pneumatron for species from two sites with different climate conditions. Two years old plants (*n =* 4) of ‘Doppio Sanguinello’ blood orange (*Citrus sinensis* (L.) Osbeck grafted on *Poncirus trifoliata* (L.) Raf. were grown under greenhouse conditions in 5 L pots filled with commercial soil (Flora-Toskana GmbH, Kempten, Germany). To identify the potential occurrence of problems in the VCs for other species, we evaluated trees from a forest near Ulm University (Germany, 48°25ʹ20ʹʹ N, 9°57ʹ20ʹʹ E, 620 m a.s.l.): *Acer campestre* (L.) (*n =* 6)*, Acer pseudoplatanus* (L.) (*n =* 3)*, Carpinus betulus* (L.) (*n =* 6)*, Fagus sylvatica* (L.) (*n =* 6)*, Populus tremula* (L.) (*n =* 4)*, Prunus avium* (L.) (*n =* 3)*, Quercus petraea* (Matt.) Liebl (*n =* 6), and *Quercus robur* (L.) (*n =* 6). For *Coffea arabica* (L.) (*n =* 3), we used potted plants growing under greenhouse conditions at Ulm University. Six years old plants of *Coffea arabica* were grown in 18 L pots filled with a standardized soil type ED 73. Branches of *Olea europaea* (L.) (*n =* 5) were collected from trees at the Agronomic Institute in Campinas SP Brazil (22°52’18” S, 47°04’39” W, 679 m a.s.l.). For *Eucalyptus camaldulensis* Dehnh (*n =* 3), we used the same curves presented previously (Pereira *et al*., 2020), obtained from plants growing in Campinas SP Brazil.

Branches were cut from plants early in the morning and transferred into plastic bags with the cut end under water to the laboratory, which took about five minutes. Branches were kept in moist plastic bags in the dark for at least 60 min to ensure that stomata were closed before preparing the branch and leaves for imaging and pneumatic measurements.

### Pneumatic measurements in branches

A Pneumatron apparatus was used to measure the gas diffusion kinetics of desiccating branches (Pereira *et al*., 2020; Jansen *et al*., 2020; Trabi *et al*., 2021). Measurements were taken every 15 minutes and the xylem water potential was monitored in bagged leaves with a pressure chamber (PMS 1505D, PMS Instruments, Corvalis OR, USA). Using a vacuum pump, 40 kPa of absolute pressure was applied to a rigid tube with known volume that was connected with the proximal end of a branch to extract gas. The vacuum pump reached 40 kPa (i.e., the initial pressure Pi) within less than a second and pressure was recorded every 500 ms during the gas discharge. The final pressure (Pf) was taken after 15 s. According to the ideal gas law, the moles of air extracted from vessels (Δn, mol) were given as follows:

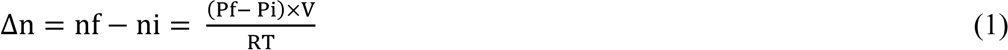

where ni and nf represented the initial and final number of moles of air at the initial and final pressure, respectively. V was the fixed discharging tube volume (in liter), R is the gas constant (8.314 kPa·L·mol^−1^·K^−1^), and T was the room temperature (25°C).

The equivalent amount of gas discharged (GD, in μL) at atmospheric pressure (P_atm_, 98 kPa) and the percentage of gas discharged (PGD, %) were calculated as:

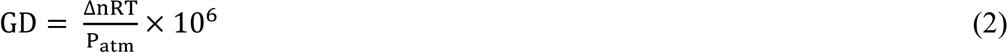

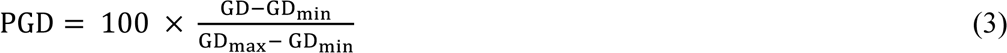

where GD_min_ was the minimum volume of air when the branch was well-hydrated, and GD_max_ was the maximum volume when the branch was severely dehydrated and GD stopped increasing even with decreasing xylem water potential.

VCs were then generated by plotting PGD or percentage of embolised pixels (PEP, as described below for the optical method) against xylem water potential (Ψ) and fitted by the following equation (Pammenter and Vander Willigen, 1998):

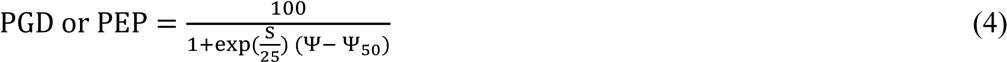

where S represented the slope of the curve, and Ψ_50_ the xylem water potential corresponding to 50% of GD_max_. The xylem water potential at 12% and 88% of gas discharged, known as Ψ_12_ and Ψ_88_ respectively, were calculated following Domec & Gartner (2001):

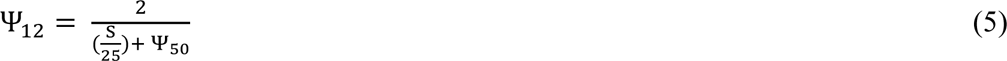

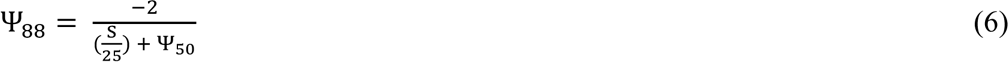

### Adjustment of the Pneumatic measurements

To determine the beginning of the plateau in the vulnerability curves obtained with the Pneumatron, the slope between two points/measurements was calculated as:

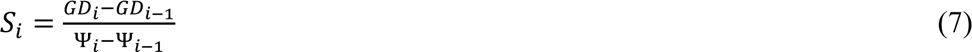

where S_i_ is the slope between two points/measurements, GD_i_ and Ψ_i_ are the volume of gas discharge and water potential at a measurement point and GD_i-1_, and Ψ_i-1_ are the volume of gas discharge and water potential one-hour prior to GD_i_ and Ψ_i_. Although pneumatic measurements were taken every 15 minutes, the slope was calculated between one point and the point one hour prior to minimize noise between measurements. By plotting the slope of the VC by each measurement point in time order (Fig. 1a,c), we were able to identify when there was a shift in GD at the beginning of the curve (Fig. 1d). When there was an initial shift in GD, the slope was higher at the beginning of the curve and then reached values close to zero, which represented the initial plateau (Fig. 1c). In that case, the initial plateau was automatically identified as the smallest slope (close to zero) at the beginning of the curve, but after seeing an initial shift in GD. In this case, all points preceding the beginning of the initial plateau were removed. When there was no initial shift in GD, the slope of the curve at the beginning of the measurements showed values close to zero, and no adjustment or correction was applied (Fig. 1a,b).

**Figure 1:**
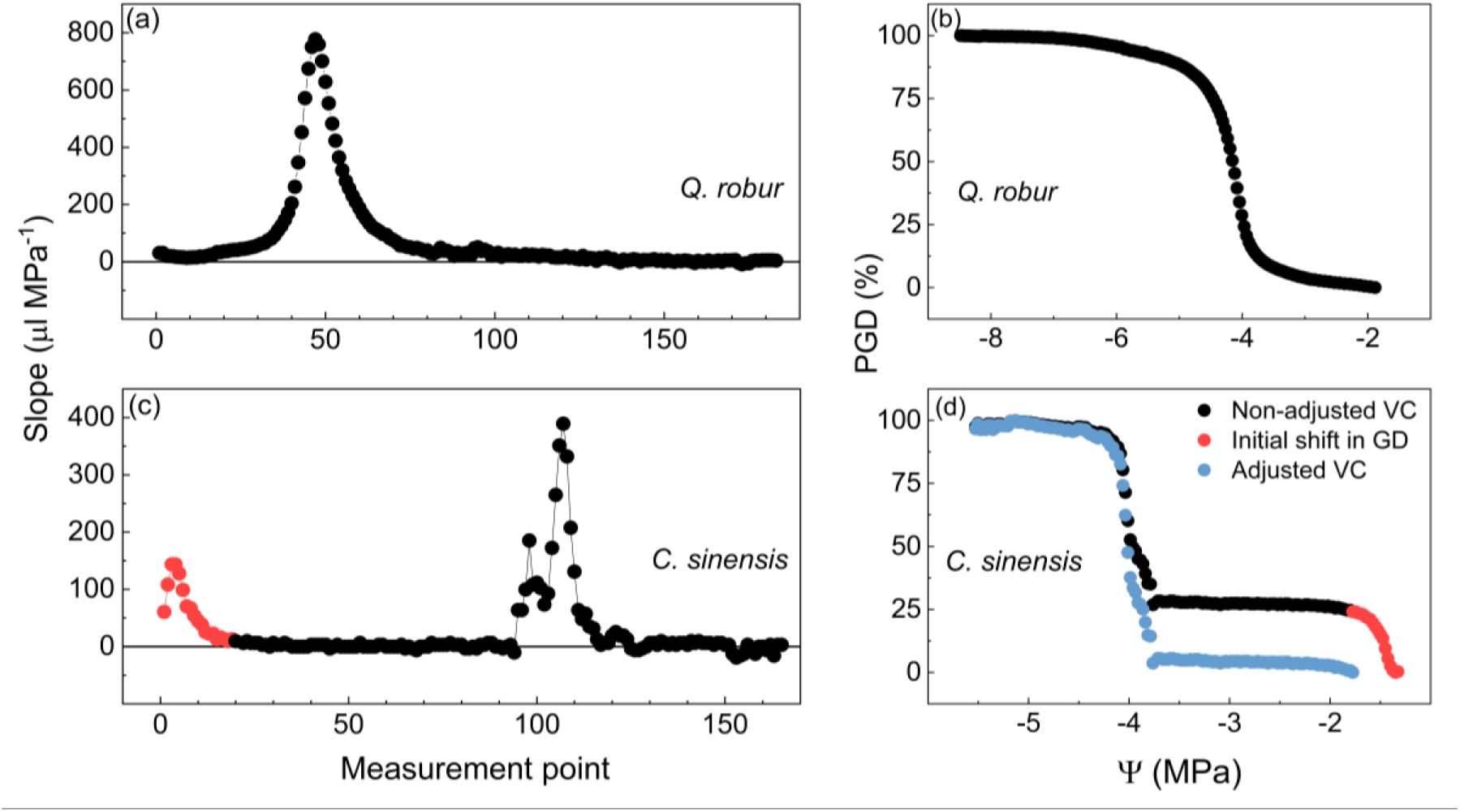
Comparison of pneumatic vulnerability curves without [*Q. robur* in (a) and (b)], and with an initial shift in gas discharge [*C. sinensis* in (c) and (d)]. First, the slope of the pneumatic vulnerability curve (VC) for each measurement point during sample dehydration was plotted [in (a) and (c)]. When there was no initial shift in gas discharged (GD), the initial slope was close to zero, providing an initial plateau in the VC (a), and no adjustment was performed (b). When there was an initial shift in GD, the slope near the dehydration start was far from zero, as shown by the red symbols in (c). In that case, the points corresponding to the initial shift in GD (red points in d) were removed and the curve was fitted again (blue symbols in d).

### Optical measurements on leaves and branches of Citrus plants

A healthy terminal *Citrus* branch of approximately 3 to 6 mm in diameter and 50 cm in length was selected and a leafless region of the branch was prepared. A region of bark approximately 15 to 20 mm in length was carefully removed from one side of the branch to expose the underlying wood without causing damage to the xylem. Once a window was created, it was firmly secured onto a stereo microscope (Axio Zoom.V16, Zeiss, Jena, Germany) using duct tape to ensure no movement of the sample while drying. A thin layer of hydrogel (Tensive Gel, Parker Laboratories, Fairfield NJ, USA) was applied to the exposed xylem surface to improve light transmission and reduce evaporation from the surface (Brodribb *et al*., 2017).

For leaves, we used both the same stereo microscope cited above and optical clamps (for more details, see http://www.opensourceov.org/) (Brodribb *et al*., 2016). A healthy, mature and undamaged leaf was selected from each branch and fixed with duct tape under the stereo microscope or in optical clamps. During the scanning, the leaf remained attached to the branch. The leaf surface area (1 cm^2^) was scanned while the leaf dried.

For both leaves and branches, images were taken every 5 min, and the water potential was monitored in bagged leaves with the pressure chamber mentioned above. For measurements of leaf water potential, we used other leaves, similar in size, on the same branch of the leaf that was being scanned and exposed to similar conditions. The water potential between each measurement interval was estimated assuming a linear decrease during dehydration (Pereira *et al*., 2020). Images were processed using ImageJ and the “OpenSourceOV ImageJ Toolbox” was used to analyse the images. Embolism events were determined by analysing differences in pixels of individual images from the previous images due to changes in the brightness of the xylem, which were then transformed into masks for quantification. The Percentage of Embolised Pixels (PEP) was quantified as the sample dried over time (Brodribb *et al*., 2016, 2017) and Ψ_50_, Ψ_12_ and Ψ_88_ were calculated using equations 4 to 6.

The Pneumatron was connected to the part of the *Citrus* branch that was cut from the whole plant, while a leaf or branch from the same terminal branch was selected for the optical measurements. For *A. pseudoplatanus, C. betulus, F. Sylvatica, P. avium, Q. petraea* and *Q. robur*, we used optical method data on embolism resistance from Guan *et al*. (2022). In this study, samples were collected at Ulm University, in the same forest from which we took some of our samples for VCs using the Pneumatic method. Therefore, were able to compare Ψ_12_, Ψ_50_ and Ψ_88_ values obtained with the Pneumatic method with those obtained by Guan *et al*. (2022) with the optical method.

### Sap extraction from cut-open vessels in Citrus

This experiment was made to measure the amount of water extracted from cut-open vessels of *Citrus* branches under negative pressure. Branches of 5 to 15 mm in diameter were cut from plants early in the morning and transferred into plastic bags to the laboratory. The transfer took about five minutes. Once in the lab, branches were let to dehydrate on the bench top for different periods. We used one branch for each measurement. After a certain amount of dehydration, the branch was put in a dark plastic bag for one hour to minimize transpiration. Then, xylem water potential was measured in a leaf using the pressure chamber mentioned earlier, and the terminal part of the branch was cut off, leaving a leafless segment of about 10 cm. We choose 10 cm segments to make sure that most of the vessels were cut open as the mean vessel length of *Citrus sinensis* plants is about 24 cm (data not shown). Immediately after cutting, the proximal end of the segment was connected to the Pneumatron apparatus and a vacuum reservoir of 100 mL using elastic tubing and plastic clamps. Inside the elastic tubing, there was an 8 mm cotton filter previously weighted (W_1_) in direct contact with the sample to collect exudated sap. A vacuum of about 40 KPa was applied for 2 minutes to the sample, and then the cotton filter was weighed again (W_2_). Then, branch dry mass (B_dry_) was determined after drying samples in an oven with forced air circulation at 60 °C until constant weight. We assumed that the weight difference between cotton filters before and after the vacuum (W_1_ and W_2_, respectively) was due to sap extracted from the cut-open vessel. Since samples were not the same size, we standardized data by dividing the difference in weight in the filter by the dry mass of each sample:

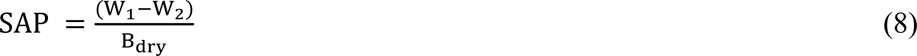

where SAP is the mass of sap extracted from samples with cut open vessel standardized by the sample dry mass (B_dry_), expressed in mg of extracted sap per g of dry mass.

### Turgor loss point

The xylem water potential at turgor loss point (Ψ_TLP_) was based on literature data for *A. campestre*, *A. pseudoplatanus*, *C. betulus*, *C. sinensis*, *C. arabica*, *E. camaldulensis*, *F. sylvatica*, *O. europaea*, *P. tremula*, *P. avium, Q. petraea*, and *Q. robur* (Supl. Table 1).

### Data analyses

Data processing and statistical analyses were performed using R v.4.3.0 (R Core Team, 2013), OriginPro v.9.3 (OriginLab Corporation, Northampton MA, USA), and JASP software (https://jasp-stats.org). Pearson’s correlation coefficient was used to test for linear correlation. Values of Ψ_12_, Ψ_50_ and Ψ_88_ were compared using Bayesian statistics. After the detection of significant effects in Bayesian ANOVA, Bayes Factors (BF_10_) were used to compare mean values. Our interpretation of Bayes Factors as evidence for alternative hypothesis (H_1_) was: 1 < BF_10_ < 3 indicates weak support for H_1_, 3 < BF_10_ < 20 indicated positive support for H_1_, while BF_10_ > 20 indicated strong support for the alternative hypothesis (Miranda *et al*., 2021).

## Results

### Embolism vulnerability curves of Citrus branches and leaves

When constructing pneumatic VCs of *Citrus* plants, some curves were exponential, with Ψ_50_ close to -1 MPa and positive values of Ψ_12_ (Fig. 2). A large discrepancy between the curves and therefore between Ψ_12_ and Ψ_50_ values was found (Fig. 2). Mean values of Ψ_12_ and Ψ_50_ were -0.9 MPa and -2.4 MPa, with a standard deviation of 1.7 and 1.5 MPa, respectively.

**Figure 2:**
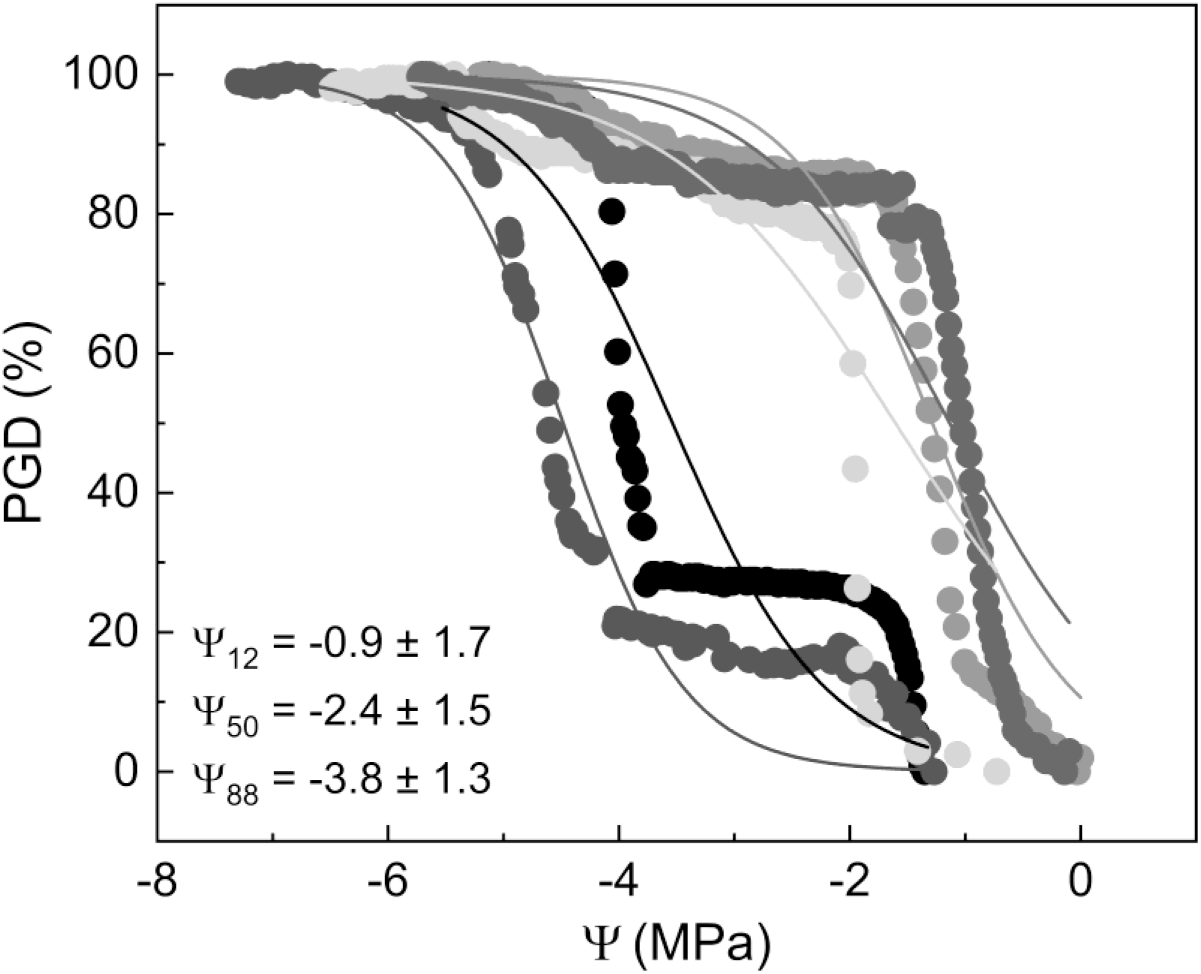
Embolism vulnerability curves of *Citrus* branches obtained with a Pneumatron showing five replicates with varying curve shapes. Lines represent the logistic function by Pammenter and Vander Willigen (1998). Ψ_12_, Ψ_50_ and Ψ_88_ mean values ± standard deviation are shown. PGD stands for the percentage gas discharge extracted with a Pneumatron, and Ψ is the xylem water potential.

Embolism resistance of leaf and branch xylem based on the optical technique revealed mean Ψ_50_ of -3.63 ± 0.2 and -4.13 ± 0.6 MPa for branches and leaves, respectively (Fig. 3). There was no statistical difference in Ψ_12_ and Ψ_50_ between leaves and branches using the optical method, while Ψ_88_ was less negative for branches (BF_10_ = 3.2).

**Figure 3:**
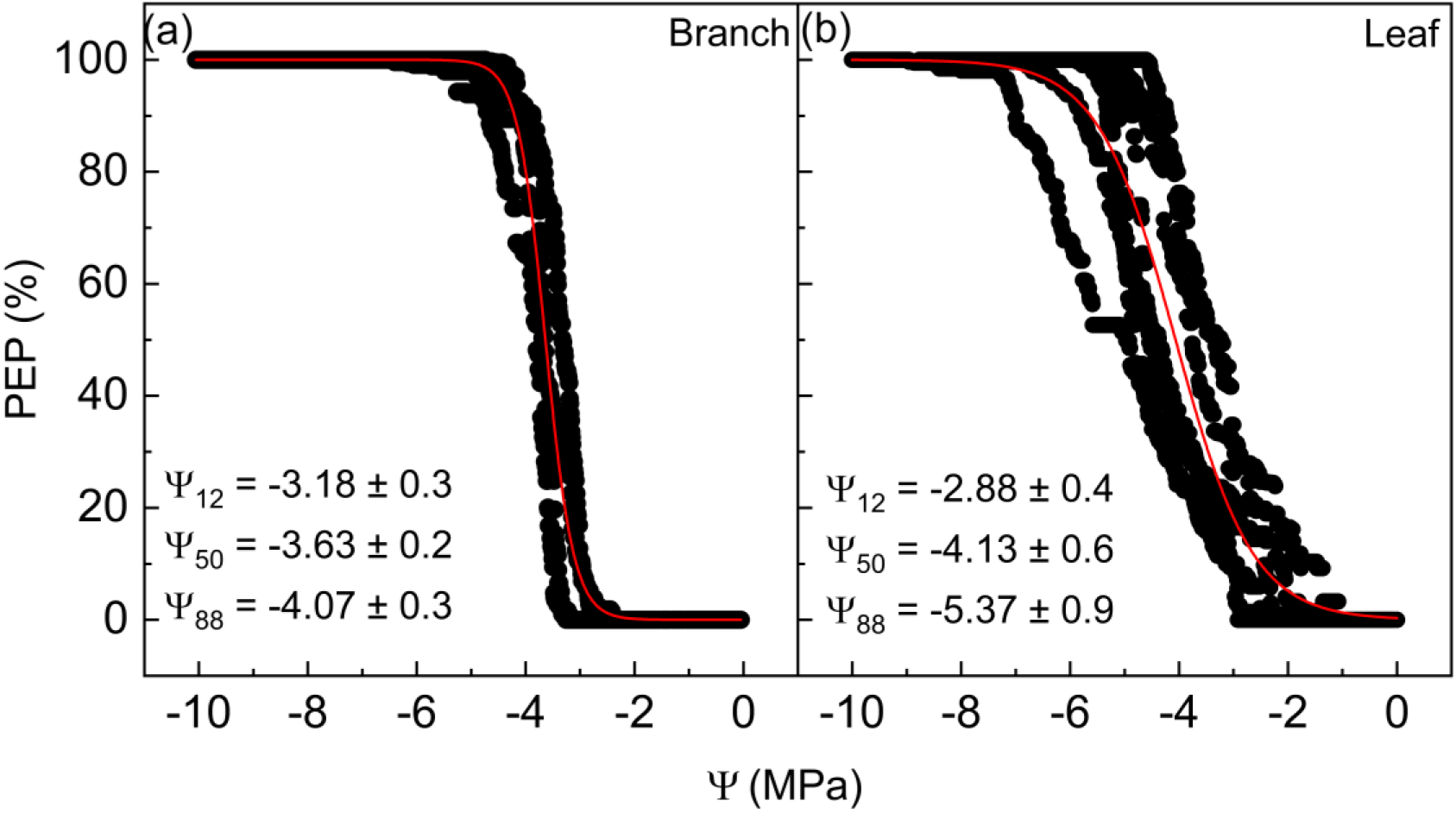
Embolism vulnerability curves of *Citrus* plants using the optical method in branches (a, n=4) and leaves (b, n=8). Red lines represent the logistic function by Pammenter and Vander Willigen (1998). Ψ_12_, Ψ_50_ and Ψ_88_ values (MPa) are presented, mean ± standard deviation. PEP stands for the percentage of embolised pixels, and Ψ is the xylem water potential.

### Sap extracted from Citrus cut-open vessels

Sap was extracted from 10 cm long *Citrus* segments by connecting the branch end to a Pneumatron. We found an exponential correlation between the sap extracted after applying the vacuum and the xylem water potential. At xylem water potentials below -1 MPa, a considerably high amount of sap was extracted from the branches (Fig. 4). Sap was no longer extracted at water potentials close to the turgor loss point for *Citrus sinensis* (-2.3 MPa)

**Figure 4:**
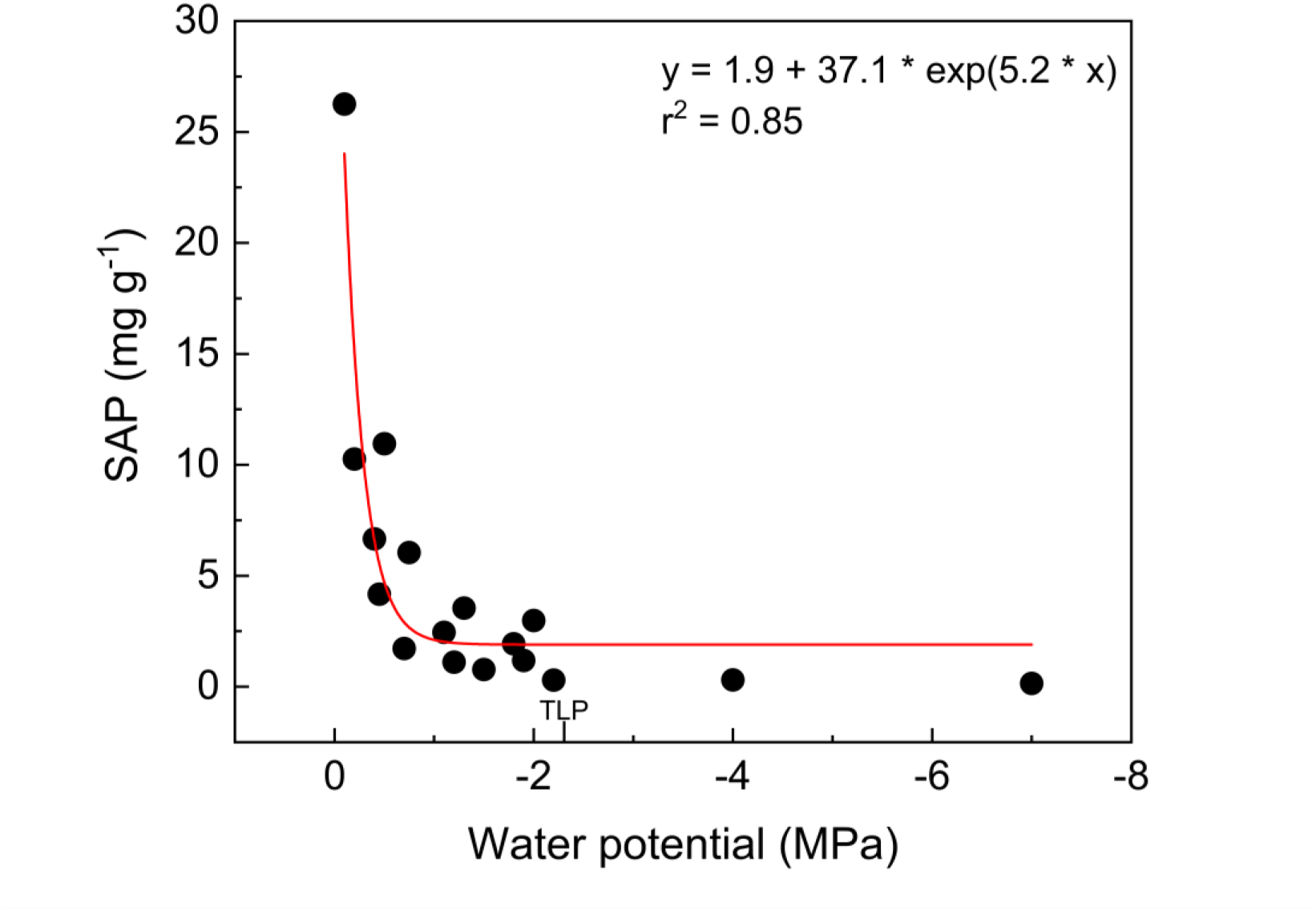
Relationship between the weight of sap extracted from cut-open vessels (SAP in mg of extracted sap per g of branch dry weight) in 10 cm long branches of *Citrus sinensis*, and xylem water potential measured during bench dehydration. Each symbol represents one measurement, and there were 17 measurements. The red line represents the exponential relationship, and the equation and r² value are shown. The mean value of the osmotic potential at the turgor loss point (TLP) is indicated on the x-axis (-2.31 MPa).

### Adjustment of VCs by considering the stability of the initial gas discharge plateau

Similar observations as for *Citrus* were observed when analysing pneumatic vulnerability curves of various species from temperate and tropical environments: an increased amount of gas discharge occurred in several but not all species. Detecting and adjusting the VCs was done by using the method described above in the Material and Methods section. Then, VCs with and without an initial increase in the amount of gas discharged were compared to each other (Fig. 5). In all adjusted VCs, the initial plateau was clearly identified before the turgor loss point (Ψ_TLP_), as shown by the dashed lines in Fig. 5. In four of the 12 studied species (*E. camaldulensis, O. europaea, Q. petraea,* and *Q. robur*) there was no need for adjustment, because the VCs showed a stable plateau during the first stages of sample dehydration, and thus no initial shift in the amount of gas extracted (Fig. 5). For eight species (*A. campestre, A. pseudoplatanus, C. betulus, C. sinensis, C. arabica, F. sylvatica, P. tremula* and *P. avium*) an early shift in GD was detected (Fig. 5). After removal of this initial change, the corrected VCs showed considerably reduced standard deviations of derived parameters Ψ_12_ and Ψ_50_ (Figs. 5, 6 and Table S2).

**Figure 5:**
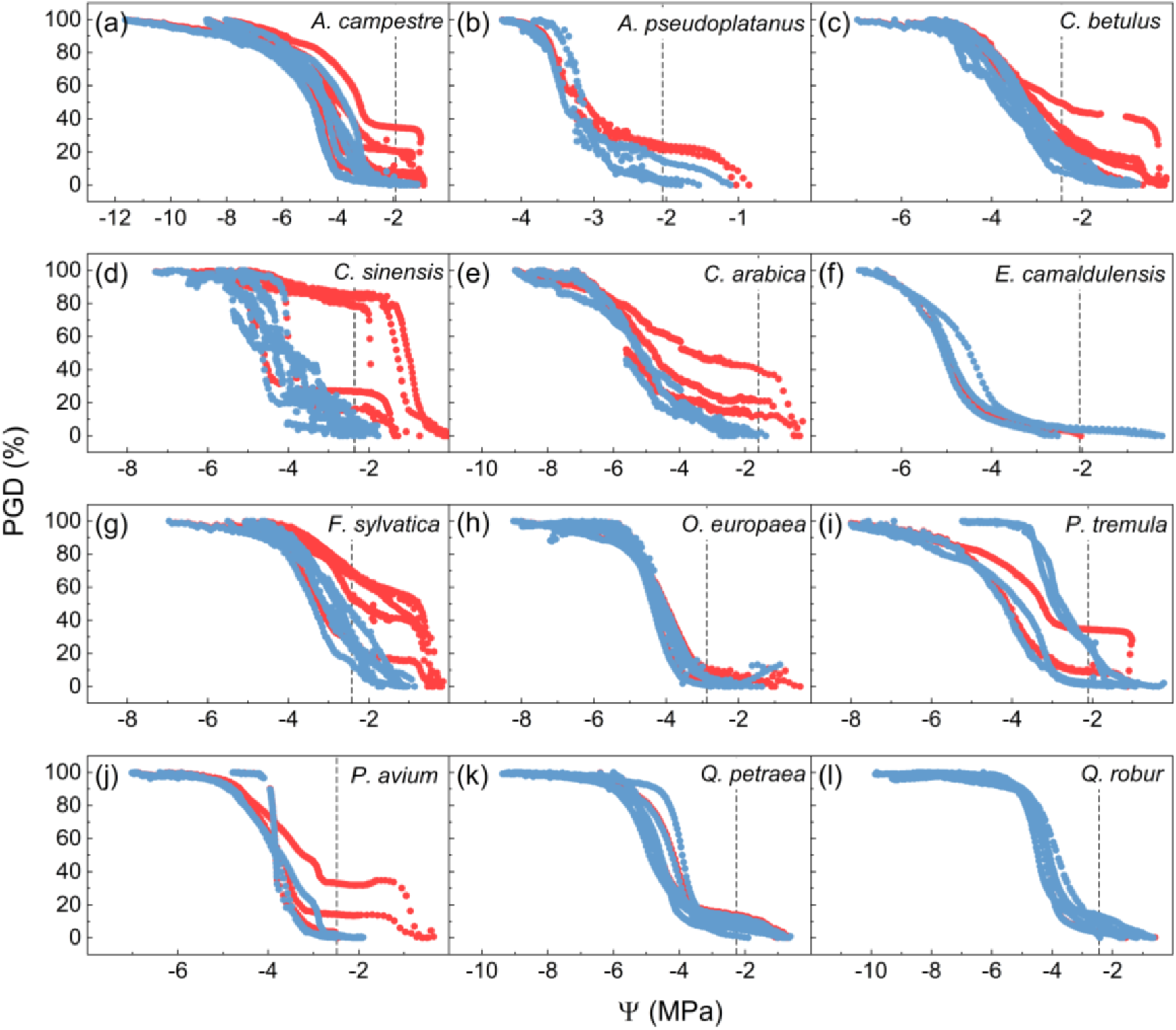
Embolism vulnerability curves obtained with a Pneumatron without (red symbols) and with (blue symbols) correction for xylem sap residue in cut-open conduits. Dashed lines indicate the water potential at the turgor loss point (Ψ_TLP_). Species included *Acer campestre* (a, n*=*6)*, Acer pseudoplatanus* (b, n*=*3)*, Carpinus betulus* (c, n*=*6)*, Citrus sinensis* (d, n*=*5), *Coffea arabica* (e, n*=*3), *Eucalyptus camaldulensis* (f, n*=*3)*, Fagus sylvatica* (g, n*=*6)*, Olea europaea* (h, n*=*5), *Populus tremula* (i, n*=*4)*, Prunus avium* (j, n*=*3)*, Quercus petraea* (k, n*=*6), and *Quercus robur* (l, n*=*6). PGD stands for the percentage gas discharge, and Ψ is the xylem water potential.

Statistical differences between Ψ_12_ and Ψ_50_ with and without the adjustment were found for *C. sinensis* (BF_10_ = 2.8 and 2.5, respectively), and *F. sylvatica* (BF_10_ = 43 and 13, respectively) (Fig. 6a,b). Besides, Ψ_88_ was not significantly affected by the initial plateau adjustment in any species. Although we found absolute differences in Ψ_12_ when comparing VCs with adjusted and non-adjusted plateaus for *A. campestre*, *A. pseudoplatanus*, *C. betulus*, *C. arabica*, *P. tremula*, and *P. avium*, there was no statistical difference for this parameter probably due to a large standard deviation without the adjustment (Fig. 6 and Table S2). In fact, the standard deviations of all VC derived parameters were considerably reduced with the adjustment for the majority of species (Table S2).

**Figure 6:**
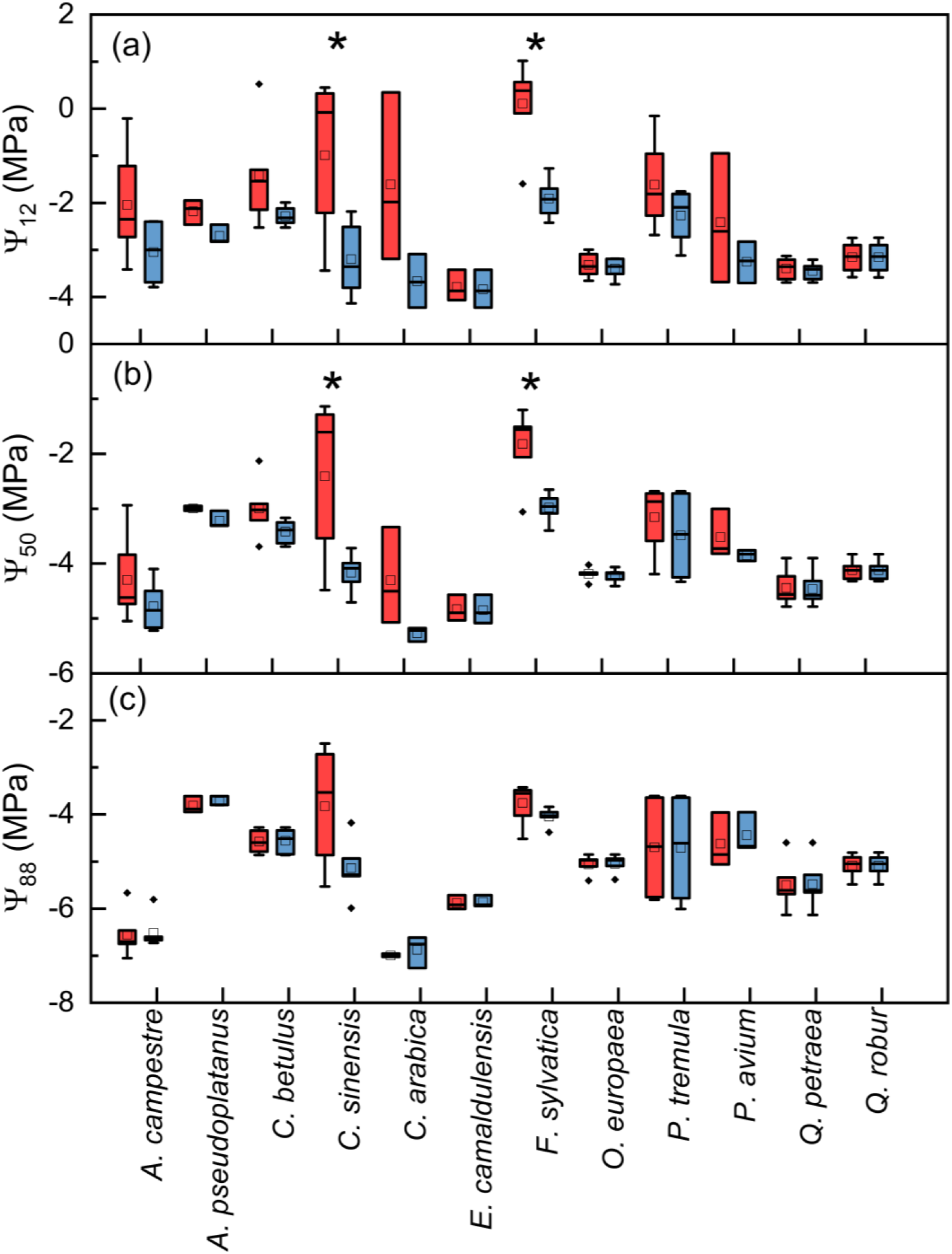
Values of Ψ_12_ (in a), Ψ_50_ (in b), and Ψ_88_ (in c) derived from VCs obtained with the Pneumatron without (red) and with (blue) correction for xylem sap residue in cut- open conduits. Boxplots consider the median, 25^th^ and 75^th^ percentiles, and mean values are represented by squares (n=3 to 6). * indicates differences between adjusted and non- adjusted values for a given species, and ◆ indicates outliers.

When comparing Ψ_12_, Ψ_50_ and Ψ_88_ values obtained with the pneumatic method with those obtained with the optical method and reported by Guan *et al*. (2022), a large discrepancy was observed for the pneumatic Ψ_12_ when no initial plateau adjustment was applied (red symbols, Fig. 7a): Ψ_12_ values were considerably higher for the pneumatic than the optical method. However, the discrepancy of Ψ_12_ and Ψ_50_ between the methods was considerably reduced when correcting the VCs (blue symbols, Fig. 7a,b).

**Figure 7:**
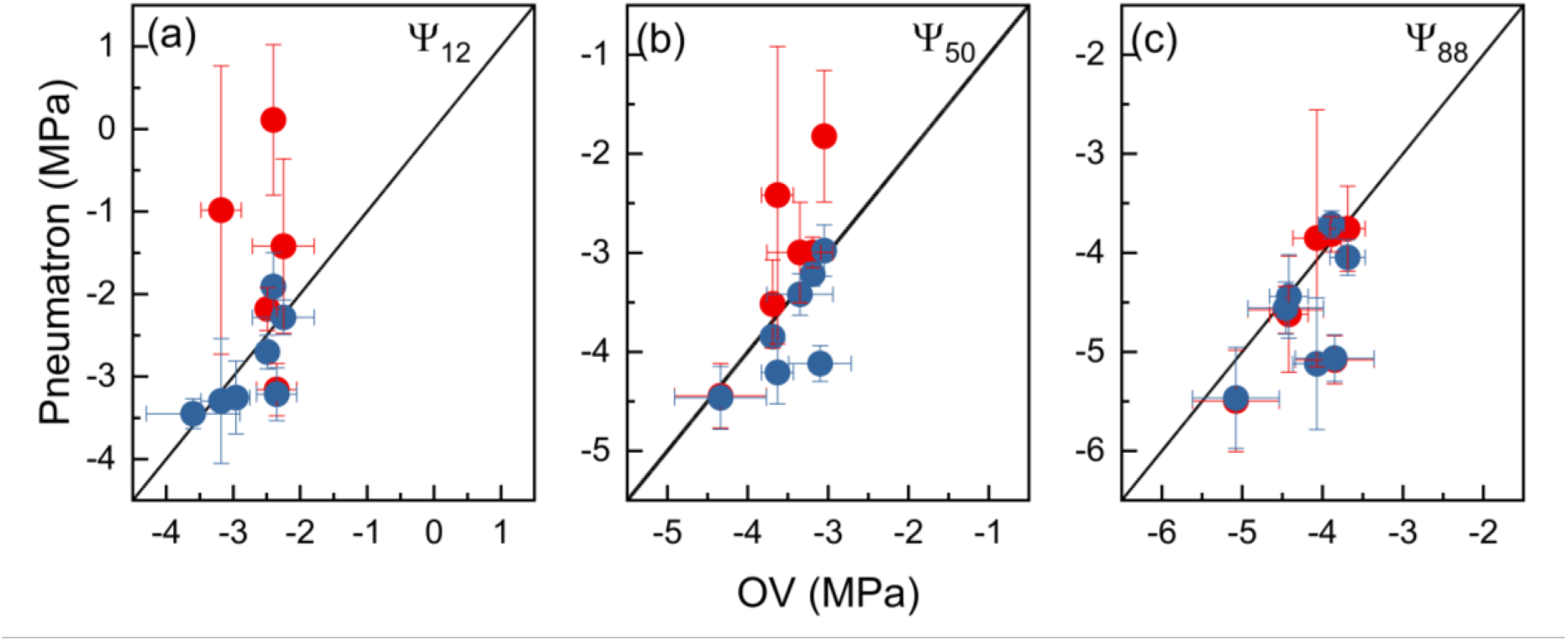
Correlations between Ψ_12_ (a), Ψ_50_ (b) and Ψ_88_ (c) obtained with the optical method (OV) and the Pneumatron method for *A. pseudoplatanus, C. betulus, C. sinensis, F. sylvatica, P. avium, Q. petraea,* and *Q. robur*. Values are presented as mean ± standard deviation. Pneumatron data were adjusted (blue) or non-adjusted (red). The OV data for *A. pseudoplatanus, C. betulus, F. sylvatica, P. avium, Q. petraea*, and *Q. robur* were taken from Guan *et al*. (2022). For *C. sinensis,* the OV data were measured and shown in Fig. 3.

## Discussion

Our experiments on *Citrus* showed that not all xylem sap in cut-open conduits may be removed when cutting well-hydrated samples in air. This observation provided an interesting explanation for changes in the gas discharge amount that do not reflect embolism propagation in intact conduits. Our sap extraction experiments also demonstrated that the amount of water that could be extracted from cut-open vessels in *Citrus sinensis* was related to the xylem water potential (Fig. 4 and 5), with hardly any sap extraction once the xylem water potential falls below the turgor loss point.

Our results also show that an increase in gas discharge during the initial stages of xylem dehydration occurred in 8 out of 12 species, although it only affected significantly the pneumatic Ψ_12_ and Ψ_50_ estimation for *Citrus sinensis* and *Fagus sylvatica* (Fig. 6). However, no effect was found for the estimation of Ψ_88_ for all species. Our analysis of the slopes of the pneumatic VCs allowed us to detect the artificial increase in gas discharge due to withdrawing xylem sap, and improved the estimation of embolism resistance. Evidence for the applied correction is that the pneumatic VCs were largely similar to the VCs obtained with the optical method (Fig. 7). In general, the absolute effect of the correction was more pronounced for Ψ_12_ than for Ψ_50_ and Ψ_88_ (Figs. 5 and 6). Besides, the standard deviation was considerably reduced when the stability of the initial plateau in VCs was corrected. The standard deviations for Ψ_12_ and Ψ_50_ of *Citrus* plants, for instance, were reduced from ±1.7 to ±0.8 MPa, and from ±1.5 to ±0.3 MPa, respectively (Fig. 6 and Table S2).

The shift in GD during initial dehydration stages was found in 8 species, and we noticed minimal differences when comparing adjusted and non-adjusted Ψ_50_ values for 10 out of the 12 species studied. The large intraspecific variability in whether or not this artifact occurs may explain why some previous methodological comparisons on embolism resistance did not detect the artefact reported here (Zhang *et al*., 2018; Sergent *et al*., 2020; Pereira *et al*., 2020a; Guan *et al*., 2021; Paligi *et al.,* In Press). Our findings may also explain why previous reports described a weak correlation for Ψ_12_ between various vulnerability curve methods (Fig. 7; Paligi *et al*., In press; Yang *et al*., 2023). As shown in Fig. 7a, the proposed adjustment can improve the accuracy of the Ψ_12_ estimation, at least when comparing the pneumatic and optical methods. In addition, our approach to focus on temporal changes in the slope of VCs also addresses the controversy on the subjective detection of the initial and final plateaus (Brum *et al*., 2023; Chen *et al*., 2023). As shown, these plateaus can be mathematically detected. Similar to the initial plateau, the final plateau was not an issue for all species studied here. We suggest to consider the final plateau within a range of xylem water potentials that is physiologically relevant for embolism formation, and not outside the measuring range of the Scholander pump or stem psychrometers.

According to the Cohesion-Tension theory, sap will be under negative pressure if the plant is transpiring at the time the conduit is cut open. Thus, xylem sap should be drained from open vessels to rehydrate the surrounding tissue immediately after cutting these (Dixon & Joly, 1895; Tyree *et al*., 2003). Our explanation for the artifact observed, however, is that xylem sap is most likely not quickly removed from cut-open conduits when the plants are well hydrated. When applying a partial vacuum with the Pneumatron, the amount of sap extracted from fully cut-open conduits in short branch segments of *Citrus* depended on the water potential. The highest amount of sap was extracted under well hydrated conditions, while sap extraction was no longer observed beyond the turgor loss point (Fig. 4). Based on this, we can assume that water in cut-open vessels was not immediately absorbed by surrounding tissues in some species. If transpiration is low and xylem water potentials are close to zero, it is possible that, after cutting xylem tissue, its sap is gradually drained by the surrounding tissue. As long as this surrounding tissue, which includes parenchyma and living fibres, has still turgor pressure, the xylem sap may not be drained immediately after cutting fresh samples (Fig. 8b). Why some species show the artefact more pronounced than others is unclear, but we speculate that variation in tissue fractions and connectivity of vessels with living cell types could be associated with the interspecific variation. When the turgor loss point is approached, a large amount of xylem sap is quickly absorbed and replaced by gas in cut-open conduits, which explains a fast increase of GD during the initial pneumatic measurements (Fig. 8d).

**Figure 8:**
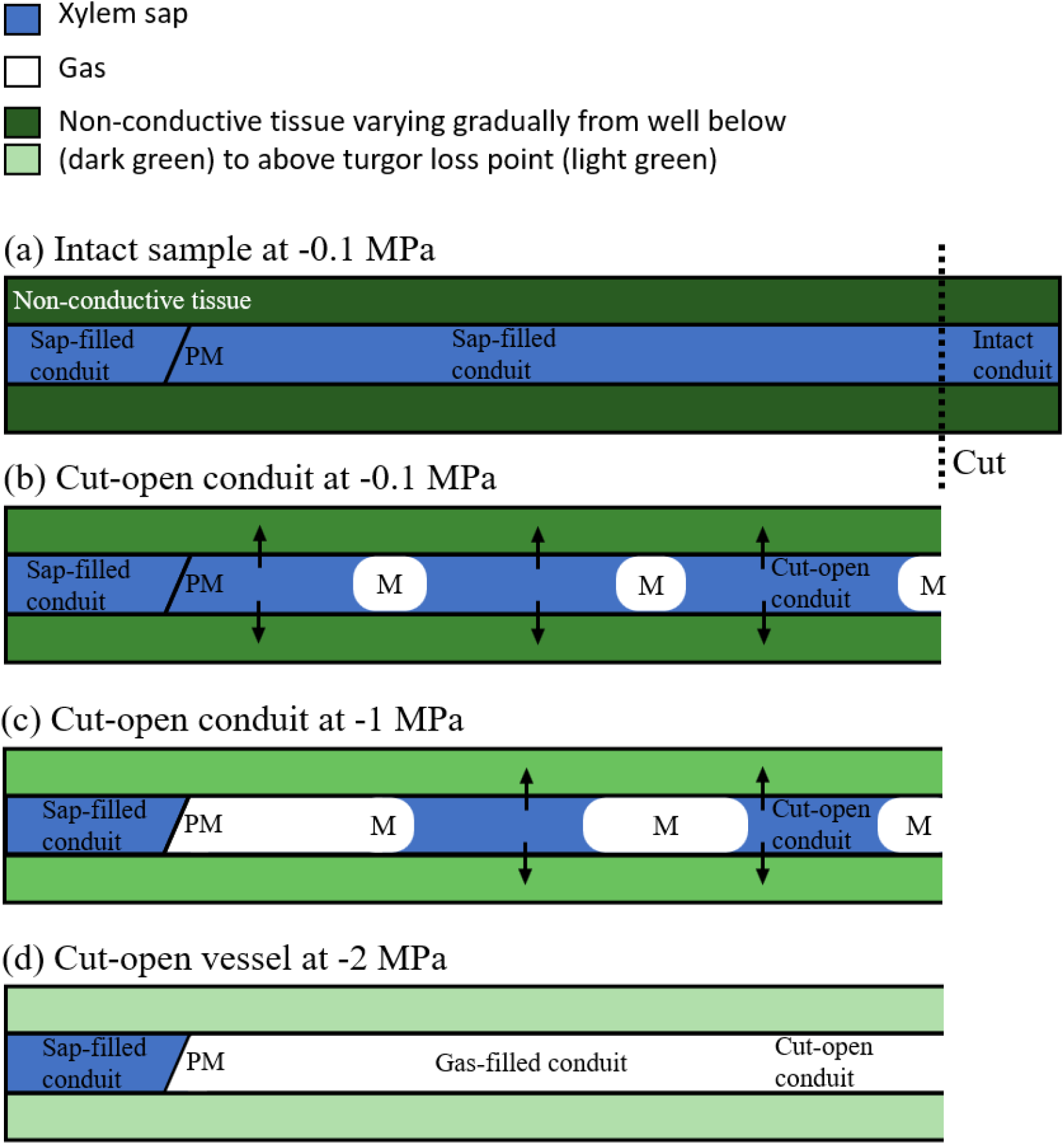
Gas-liquid dynamics in a hypothetical branch with a turgor loss point of -2 MPa. (a) hydrated intact branch, before cutting it at -0.1 MPa, with sap-filled conduits. (b) the same branch at high water potential after cutting, some menisci are formed as large gas bubbles, and water starts to be drained to rehydrate the surrounding tissue. As this tissue still has turgor (given by the dark green colour in surrounding, non-conductive tissue), not all sap is drained immediately. (c) more xylem sap is absorbed by the non- conductive cells as surrounding tissues dehydrate further. (d) complete uptake of xylem sap by non-conductive cells, with the cut-open conduit being completely filled with air, which happens at a water potential close to the turgor loss point. PM = intervessel wall with pit membranes, and M = meniscus.

The vessels in angiosperms are commonly surrounded by highly specialized living parenchyma cells. These vessel-associated cells can be connected to vessels via half- bordered pits (Morris *et al*., 2018). Interestingly, the influx of water from vessel-associated parenchyma cells has been observed based on microCT observations, and proposed as a mechanism to refill embolised vessels (Brodersen *et al*., 2010, 2018). Knipfer *et al*. (2016), for instance, found that in excised branches, water droplets emerge from the inner vessel walls adjacent to xylem parenchyma or fibres. Thus, an alternative explanation for the water extracted from branches with cut-open vessels in our study could be the active secretion of liquid from living cells that are directly connected to conduits. For the moment, we cannot exclude this hypothesis as more observations would be needed. Vessel-associated parenchyma cells are also related to tylosis formation and gel production, where balloon-like sacs of cytoplasm enter the embolised vessel resulting in partial or full vessel occlusion (Morris *et al*., 2018). Vessel occlusion by tyloses would strongly affect gas extraction on a permanent basis. However, we can exclude this hypothesis, because the artefact described only occurred during initial dehydration stages and was not permanent.

Since it is unclear how fast xylem sap is withdrawn from cut-open conduits in a particular sample, we suggest removing all data before an initial plateau in GD has been achieved as a general protocol for pneumatic VCs, and especially if an initial increase in GD is detected. For this reason, we strongly recommend users to work with the automated Pneumatron device as the high temporal resolution will be highly appropriate for this purpose. In fact, a large similarity was found in estimations of VC-derived parameters obtained with the pneumatic and optical methods when there was no initial increase in GD (Figs. 7). Also, the adjustment suggested can be useful to reduce the variability in embolism resistance, even if the artificial increase in GD is small. For instance, applying a correction to data obtained for *C. arabica* reduced the standard deviation from ±1.8 to ±0.6 MPa, and from ±0.9 to ±0.1 MPa for Ψ_12_ and Ψ_50_, respectively (Fig. 6 and Table S2).

Removing GD measurements before an initial plateau has been reached is a reasonable approach for improving the estimation of Ψ_12_ and Ψ_50_ in some species, since xylem sap appears to be completely absorbed when the surrounding tissues lose turgor. Yet, users will not be required to accurately measure Ψ_TLP_, because a stable plateau in GD should be obtained before the turgor loss point has been reached. The removal of all points before the initial plateau in pneumatic VCs would still be a safe strategy since embolism formation is unlikely before plants lose cell turgor. Previous studies have shown that stomata close at or before turgor loss, earlier than the onset of embolism (Brodribb & Holbrook, 2003; Nardini *et al*., 2003; Brodribb *et al*., 2016; Skelton *et al*., 2018). In addition, Martin-StPaul *et al*. (2017) found that most species close their stomata at water potentials long before embolism starts.

A practical consequence of our results is that VCs obtained with the Pneumatic method can be started with slightly dehydrated branches, at least up to Ψ_TLP_, which facilitates branch sampling when it is not possible to find a completely hydrated plant in the field, or when plants are already transpiring during the day. Starting with slightly dehydrated samples and not putting them under water after cutting would help the gas-liquid meniscus to pull back more quickly, which would especially be appropriate for drought-resistant species with fairly high embolism resistance, such as *Citrus* species.

## Conclusion

For the first time, we report the presence of xylem sap in cut-open vessels affecting embolism vulnerability curves obtained with the pneumatic method. As an easy correction for improving the estimation of important parameters related to embolism resistance, we propose the removal of all points before the initial plateau in pneumatic VCs if an initial shift in GD is detected based on a temporal analysis of the VC slope values. As long as a Pneumatron with high temporal resolution is used, it is feasible to detect a stable plateau before embolism occurs.

## Acknowledgments

The authors acknowledge the São Paulo Research Foundation (FAPESP, Brazil; Grant #2018/09834-5, #2019/15276-8, and #2021/13329-7), the National Council for Scientific and Technological Development (CNPq, Brazil; Grant #303664/2020-7, #311345/2019-0 and #304295/2022-1), and the Deutsche Forschungsgemeinschaft (DFG, Germany, project 508216003) for support received.

## Author contributions

MTM, LP, SJ, ECM and RVR developed the hypotheses and planned the experiments which were conducted by MTM, LP, GSP, XG, LMS and SK. MTM, LP and GSP analyzed the data. All authors contributed to the hypotheses’ discussion and manuscript writing, with substantial inputs from MTM, LP, SJ and RVR.

## Supporting Information

Table S1. The xylem water potential at turgor loss point for the species studied.

Table S2: Variation of Ψ_12_, Ψ_50_ and Ψ_88_ obtained from VCs obtained with the Pneumatron.

