## Supporting Information for "Xylem sap residue in cut-open conduits can affect gas discharge in pneumatic experiments"

**Table S1.** The xylem water potential at turgor loss point (**Ψ_TLP_**, MPa) for the species studied; average values are provided (±SD), as well as the lowest and highest values. Data were retrieved from the literature.

| **Species** | **Ψ_TLP_**  (mean and range) | **References** |
| --- | --- | --- |
| *Acer campestre* | -1.92 ± 0.40 MPa (-1.28 to -2.51) | Burghard & Riederer (2003); Nardini *et al.* (2012); Banks *et al.* (2019); Kunert & Tomaskova (2020); Thom *et al.* (2022) |
| *Acer pseudoplatanus* | -2.05 ± 0.42 MPa (-1.40 to -2.75) | Nardini *et al.* (2012); Sjöman *et al.* (2015); Li *et al.* (2016); Lübbe *et al.* (2017); Leuschner *et al.* (2019); Kunert *et al.* (2020) |
| *Carpinus betulus* | -2.45 ± 0.35 MPa (-1.85 to -2.71) | Li *et al.* (2016); Lübbe *et al.* (2017); Leuschner *et al.* (2019); Kunert *et al.* (2020) |
| *Citrus sinensis* | -2.31 ± 0.49 MPa (-1.69 to -2.82) | Savé *et al*. (1995); Gonçalves *et al.* (2016); Silva *et al*. (2019); Miranda *et al*. (2022); |
| *Coffea arabica* | -1.60 ± 0.20 MPa  (-1.34 to -1.82) | Meinzer *et al.* (1990); Da Matta *et al.* (1993); Nardini *et al.* (2014) |
| *Eucalyptus camaldulensis* | -2.06 ± 0.22 MPa  (-1.8 to -2.4) | Dreyer *et al.* (1992); White *et al.* (2000); Lemcoff *et al.* (2002); Siddiqui *et al.* (2008); |
| *Fagus sylvatica* | -2.41 ± 0.30 MPa  (-1.80 to -2.82) | Aranda *et al.* (1996); Backes & Leuschner (2000); Leuschner *et al*., (2001); Burghardt *et al.* (2003); Lübbe *et al.*(2017); Tomasella *et al.* (2018); Leuschner *et al.* (2019); Kunert *et al*., (2020) |
| *Olea europaea* | -2.88 ± 0.72 MPa  (-1.67 to 3.57) | Hinckley et al., (1980); Dichio *et al.* (1997); Dichio *et al.* (2003); Bacelar et al. (2006) |
| *Populus tremula* | -2.10 MPa | Kunert & Tomaskova, (2020) |
| *Prunus avium* | -2.48 ± 0.18 MPa  (-2.24 to -2.71) | Peschiutta *et al.*, (2013); Kunert *et al.*, (2020) |
| *Quercus petraea* | -2.27 ± 0.45 MPa  (-1.3 to -2.77) | Aranda *et al.*, (1996); Backes & Leuschner (2000); Tomas *et al.*, (2000); Burghardt *et al.*, (2003); Nardini *et al.*, (2012); Kunert *et al.*, (2020) |
| *Quercus robur* | -2.45 ± 0.22 MPa  (-2.16 to -2.68) | Tomas *et al.*, (2000); Kunert *et al.*, (2020); Hanley *et al.*, (2021) |

**Table S2:** Variation of Ψ_12_, Ψ_50_ and Ψ_88_ (in MPa) obtained from VCs obtained with the Pneumatron without and with adjustment based on **the initial plateau in AD**.

| Species | Ψ_12_ | | Ψ_50_ | | Ψ_88_ | |
| --- | --- | --- | --- | --- | --- | --- |
|  | non-adjusted | adjusted | non-adjusted | adjusted | non-adjusted | adjusted |
| *A. campestre* | -2.0 ± 1.1 | -3.0 ± 0.6 | -4.3 ± 0.8 | -4.8 ± 0.4 | -6.6 ± 0.5 | -6.5 ± 0.3 |
| *A. pseudoplatanus* | -2.2 ± 0.3 | -2.7 ± 0.2 | -3.0 ± 0.1 | -3.2 ± 0.2 | -3.8 ± 0.2 | -3.7 ± 0.1 |
| *C. betulus* | -1.4 ± 1.1 | -2.3 ± 0.2 | -3.0 ± 0.5 | -3.4 ± 0.2 | -4.6 ± 0.2 | -4.6 ± 0.3 |
| *C. sinensis* | -1.0 ± 1.8 | -3.3 ± 0.8* | -2.4 ± 1.5 | -4.2 ± 0.3* | -3.9 ± 1.3 | -5.1 ± 0.7 |
| *C. arabica* | -1.6 ± 1.8 | -3.7 ± 0.6 | -4.3 ± 0.9 | -5.3 ± 0.1 | -7.0 ± 0.1 | -6.9 ± 0.3 |
| *E. camaldulensis* | -3.8 ± 0.3 | -3.8 ± 0.4 | -4.8 ± 0.2 | -4.8 ± 0.3 | -5.9 ± 0.2 | -5.9 ± 0.1 |
| *F. sylvatica* | 0.1 ± 0.9 | -1.9 ± 0.4* | -1.8 ± 0.7 | -3.0 ± 0.3* | -3.8 ± 0.4 | -4.0 ± 0.2 |
| *O. europaea* | -3.3 ± 0.3 | -3.4 ± 0.2 | -4.2 ± 0.1 | -4.2 ± 0.1 | -5.1 ± 0.2 | -5.0 ± 0.2 |
| *P. tremula* | -1.6 ± 1.1 | -2.3 ± 0.6 | -3.2 ± 0.7 | -3.5 ± 0.9 | -4.7 ± 1.2 | -4.7 ± 1.2 |
| *P. avium* | -2.4 ± 1.4 | -3.3 ± 0.4 | -3.5 ± 0.4 | -3.8 ± 0.1 | -4.6 ± 0.6 | -4.4 ± 0.4 |
| *Q. petraea* | -3.4 ± 0.2 | -3.4 ± 0.2 | -4.4 ± 0.3 | -4.5 ± 0.3 | -5.5 ± 0.5 | -5.5 ± 0.5 |
| *Q. robur* | -3.2 ± 0.3 | -3.2 ± 0.3 | -4.1 ± 0.2 | -4.1 ± 0.2 | -5.1 ± 0.2 | -5.1 ± 0.2 |
